# Determining clinically relevant features in cytometry data using persistent homology

**DOI:** 10.1101/2021.04.26.441473

**Authors:** Soham Mukherjee, Darren Wethington, Tamal K. Dey, Jayajit Das

**Author notes:** These authors contributed equally to this work (Soham Mukherjee, Darren Wethington).

## Abstract

Cytometry experiments yield high-dimensional point cloud data that is difficult to interpret manually. Boolean gating techniques coupled with comparisons of relative abundances of cellular subsets is the current standard for cytometry data analysis. However, this approach is unable to capture more subtle topological features hidden in data, especially if those features are further masked by data transforms or significant batch effects or donor-to-donor variations in clinical data. We present that persistent homology, a mathematical structure that summarizes the topological features, can distinguish different sources of data, such as from groups of healthy donors or patients, effectively. Analysis of publicly available cytometry data describing non-naïve CD8+ T cells in COVID-19 patients and healthy controls shows that systematic structural differences exist between single cell protein expressions in COVID-19 patients and healthy controls.

Our method identifies proteins of interest by a decision-tree based classifier and passes them to a kernel-density estimator (KDE) for sampling points from the density distribution. We then compute persistence diagrams from these sampled points. The resulting persistence diagrams identify regions in cytometry datasets of varying density and identify protruded structures such as ‘elbows’. We compute Wasserstein distances between these persistence diagrams for random pairs of healthy controls and COVID-19 patients and find that systematic structural differences exist between COVID-19 patients and healthy controls in the expression data for T-bet, Eomes, and Ki-67. Further analysis shows that expression of T-bet and Eomes are significantly downregulated in COVID-19 patient non-naïve CD8+ T cells compared to healthy controls. This counter-intuitive finding may indicate that canonical effector CD8+ T cells are less prevalent in COVID-19 patients than healthy controls. This method is applicable to any cytometry dataset for discovering novel insights through *topological data analysis* which may be difficult to ascertain otherwise with a standard gating strategy or in the presence of large batch effects.

**Author summary:** Identifying differences between cytometry data seen as a point cloud can be complicated by random variations in data collection and data sources. We apply *persistent homology* used in *topological data analysis* to describe the shape and structure of the data representing immune cells in healthy donors and COVID-19 patients. By looking at how the shape and structure differ between healthy donors and COVID-19 patients, we are able to definitively conclude how these groups differ despite random variations in the data. Furthermore, these results are novel in their ability to capture shape and structure of cytometry data, something not described by other analyses.

## 1 Introduction

Cytometry data contain information about the abundance of proteins in single cells and are widely used to determine mechanisms and biomarkers that underlie infectious diseases and cancer. Recent advances in flow and mass cytometry techniques enable measurement of abundances of over 40 proteins in a single cell [1, 2]. Thus, in the space spanned by protein abundance values measured in cytometry experiments, a cytometry dataset is represented by a point cloud composed of thousands of points where each point corresponds to a single cell. Abundances of proteins or chemically modified forms (e.g., phosphorylated forms) of proteins in single immune cells change due to infection of the host by pathogens (e.g., a virus) or due to the presence of tumors which usually result in changes in the ‘shape’ of point cloud data measured in cytometry experiments [3–5]. Cytometry data analysis techniques commonly rely on Boolean gating and calculation of relative proportions of resulting populations as a method to compare datasets across control/healthy and experimental/diseased conditions. In recent years, state-of-the-art analyses based on sophisticated machine learning algorithms capable of mitigating batch effects, ad hoc gating assumptions, and donor-donor variability have been developed [6, 7]. However, these methods are not designed to quantitatively characterize shape features (e.g., connected clusters, cycles) in high dimensional cytometry datasets that can contain valuable information regarding unique co-dependencies of specific proteins in diseased individuals compared to healthy subjects.

Topological Data Analysis (TDA) aims to capture the underlying shape of a given dataset by describing its topological properties. Unlike geometry, topological features (e.g., the hole in a doughnut) are invariant under continuous deformation such as rotation, bending, twisting but not tearing and gluing. One of the tools by which TDA describes topological features latent in data is persistent homology [8, 9]. For example, for a point cloud data, persistent homology captures the birth and death of topological features (e.g., ‘holes’) in a dataset after building a scaffold called a simplicial complex out of the input points. This exercise provides details regarding topological features that ‘persist’ over a range of scale and thus contain information regarding the shape topology at different length scales (see Fig S1 for details). Persistent homology has been applied to characterize shapes and shape-function relationships in a wide variety of biological systems including skin pattern formation in zebra fish [10], protein structure, and pattern of neuronal firing in mouse hippocampus [11]. TDA has additionally previously been applied to identify immune parameters associated with transplant complications for patients undergoing allogenic stem cell transplant using populations of immune cell types assayed via mass cytometry [12]. However, this work did not use persistent homology or expression levels of proteins in their analysis, leaving the shape of cytometry data uncharacterized. Another work focuses on the use of TDA as a data reduction method for single-cell RNA sequencing data [13], but again do not attempt to characterize how topologies derived from point clouds differ among disparate data sources such as healthy and diseased individuals.

The challenges of directly applying current persistence methodologies to cytometry data to characterize distinguishing features between healthy and diseased states are the following: 1. Features that separate healthy from diseased state can pertain to the change in density of points in a region in point cloud data - therefore, the information of local density should be incorporated in persistent homology methods, in particular in the filtration step that brings in sequentially the simplices connecting the points. In commonly used Rips filtration [14] the density of points is not included. 2. There can be shape changes giving a different length scale in the point cloud data, such as formation of an elbow, in a diseased condition. 3. There can be systematic differences between healthy and diseased states across batch effects and donor-donor variations. Topological features should capture these global differences being oblivious to the local variations caused by measurement noise.

We address the above challenges by developing an appropriate filtration function to compute persistence and applying the method to characterize distinguishing features of non-naïve CD8+ T cells between healthy and SARS-CoV-2 infected patients.

## 2 Results

### 2.1 Persistence framework for SARS-CoV-2 infection

Topological signatures given by persistence are stable, global, scale invariant and show resilience to local perturbations [15]. It is this property of persistent homology that motivates us to use TDA in distinguishing clinically relevant features in flow cytometry data in COVID-19 patients.

#### Persistent Homology

Persistent homology builds on an algebraic structure called homology groups graded by its dimension *i* and denoted by H_i_. Intuitively, they describe the shape of the data by ‘connectivity’ at different levels. For example, H_0_ describes the number of connected components, H_1_ describes the number of holes, and, H_2_ describes the number of enclosed voids apparently present in the shape that the dataset represents. Three and higher dimensional homology groups capture analogous higher (≥ 3) dimensional features. A point cloud data (henceforth abbreviated as PCD) itself does not have much of a ‘connected structure’. So, a scaffold called a *simplicial complex* is built on top of it. This simplicial complex, in general, is made out of simplices of various dimensions such as vertices, edges, triangles, tetrahedra, and other higher dimensional analogues. Given a growing sequence of such complexes called *filtrations*, a persistence algorithm tracks information regarding the homology groups across this sequence. In our case, these complexes can be restricted only to vertices and edges. With the restriction that both vertices of an edge appear before the edge, we get a nested sequence of graphs

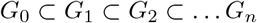

as the filtration. Fig 1 shows such a *filtration*.

**Figure 1.**
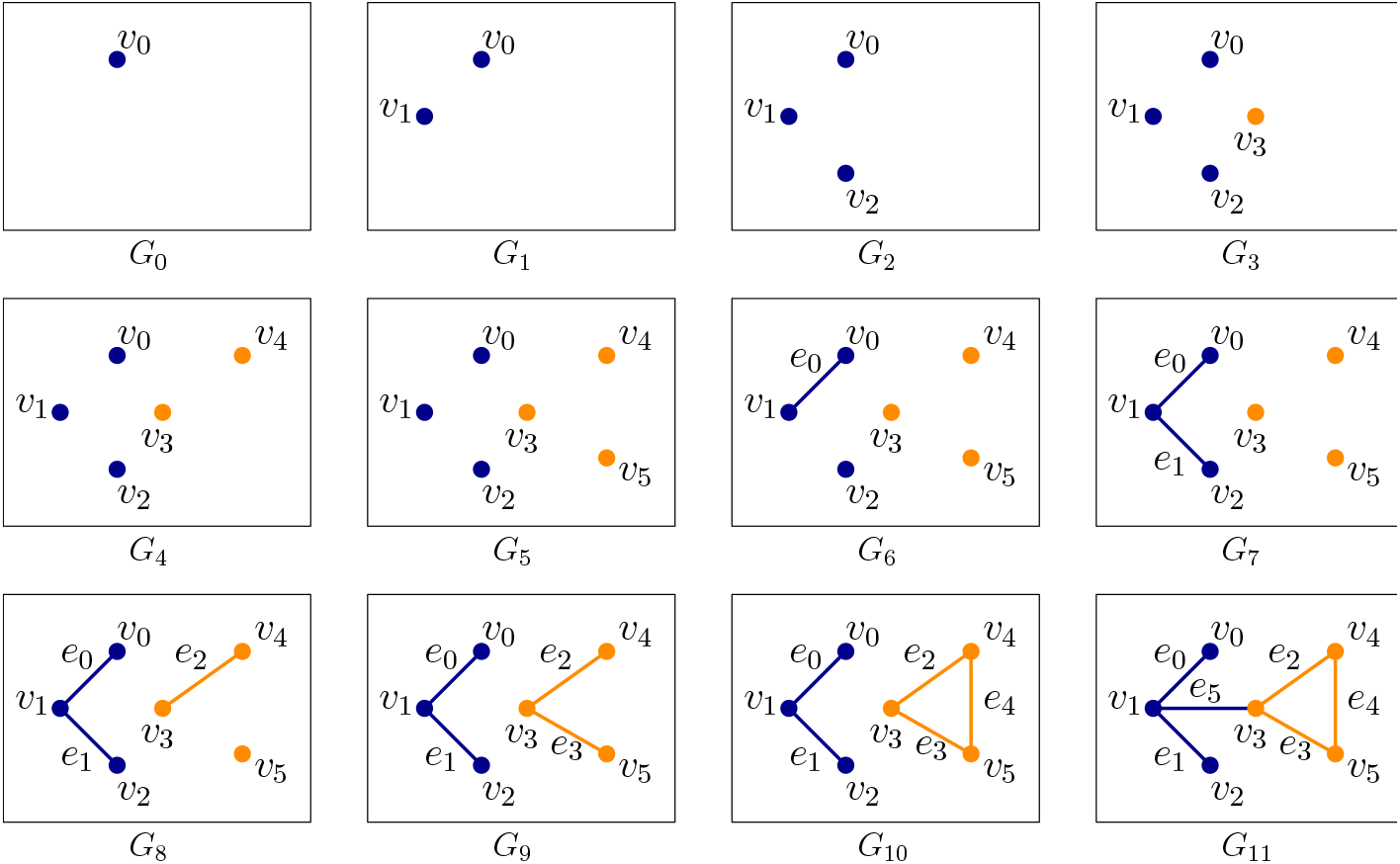
An example of *filtration* for a graph. The nested sequence of graphs *G*_0_ ⊂ *G*_1_ ⊂ … *G*_11_ forms a filtration of the final graph *G*_11_. Each vertex *v_i_* creates a new component in the nested sequence, and edges *e*_0_, *e*_1_, *e*_2_, *e*_5_ merge two components whereas *e*_4_ creates a cycle (yellow).

#### Persistence Diagram

Appearance (“birth”) and disappearance (“death”) of topological features, that is, cycles whose classes constitute the homology groups, can be captured by persistence algorithms [8, 16]. These “birth” and “death” events are represented as points in the so-called *persistence diagram*. If a topological feature is born at filtration step *b* and dies at step *d*, we represent this by *persistence pair* (*b, d*) with persistence *d* – *b*. The pair (*b*, *d*) becomes a point in the persistence diagram with the “birth” as x-axis and “death” as y-axis. This 2D plot summarizes topological features latent in the data. In the example-filtration of Figure 1 a new component gets ‘born’ when a vertex *v_i_* appears in the filtration for the first time. When an edge is introduced, one of the two things can happen–either two components are joined, or a cycle is created. In the first case, a ‘death’ happens for 0-th homology group H_0_, and in the second case, a ‘birth’ happens for the 1-st homology group H_1_. For example, when *e*_0_ comes in the filtration (*G*_6_), it merges two components created by *v*_0_ and *v*_1_. By convention, we choose to kill the component that got created later in the filtration and thus we let the component created by *v*_1_ die. We obtain a persistence pairing (1, 6) since edge *e*_0_ at filtration step 6 kills the component created by *v*_1_ at step 1. Similarly, we obtain pairs (2, 7), (4, 8), (5, 9), and (3,11). These points, tracking the ‘birth’ and ‘death’ of components, produce the persistence diagram for the 0-th homology group H_0_ and hence we refer to it as H_0_-persistence diagram. Note that the edge *e*_4_ creates a cycle (yellow) that never dies. In such cases, *i.e*. when a topological feature never dies, we pair it with ∞. For the edge *e*_4_, we obtain a persistence pair (10, ∞). But, this feature concerns the 1-st homology group H_1_ and thus it becomes a point in the persistence diagram for H_1_ which we refer to as H 1-persistence diagram. One way to leverage the above framework for studying a function is to assign function values to vertices and edges and construct a filtration by ordering them according to these assigned values. For such cases the persistence pairs take the form (*b, d*) where *b* is the value at which a feature is born and d is the value at which it dies. The function values that induce the filtration (Figure 2) are chosen to capture two features of the input PCD–(i) the density variations, and (ii) the *anisotropy* of the features, that is, how elongated it is in a certain direction, henceforth termed as *length scale* ‘of the feature’ or collectively ‘of the data’. In particular, length scales refer to the prominence of protrusions such as ‘elbows’ in COVID-19 data.

**Figure 2.**
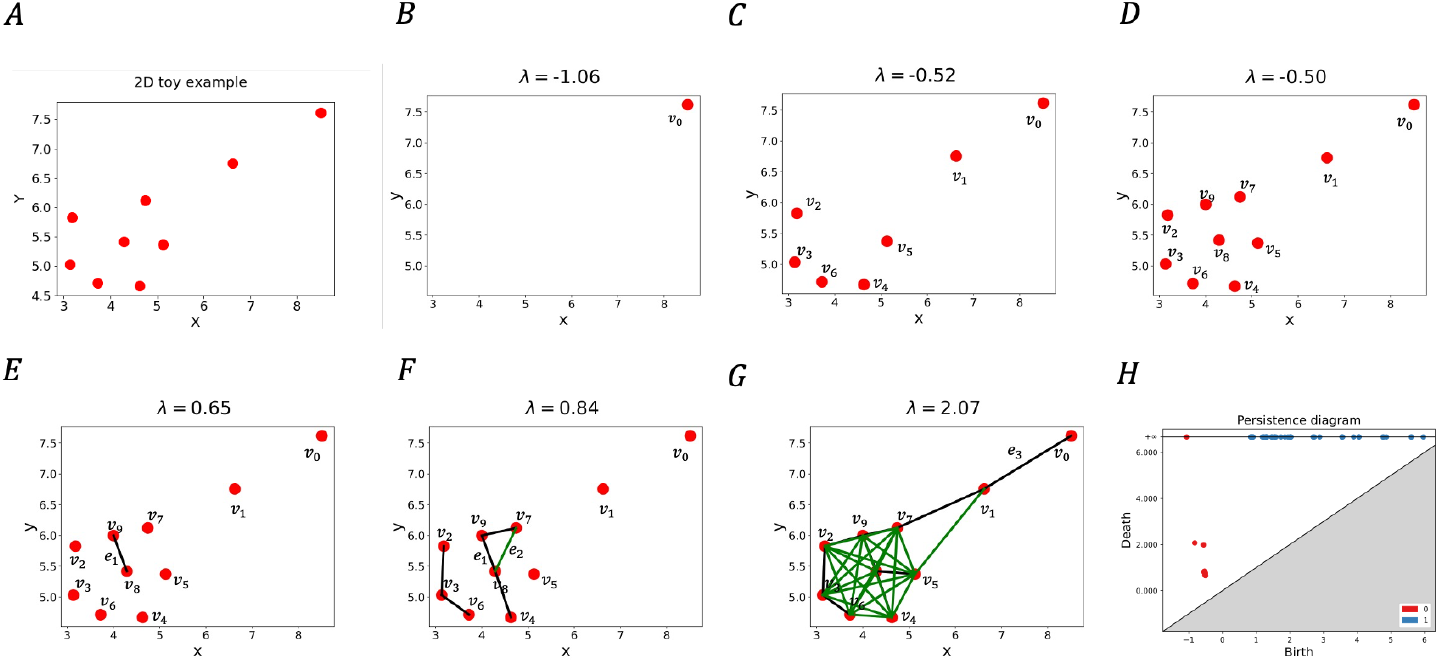
Illustration of persistence for a 2D point cloud data (PCD). **(A), (H)** shows a 2D PCD example and its computed persistence diagram. **(B)-(G)** shows important changes in topological feature as λ increases from −∞ to ∞. **(B)** At λ = −1.06 an isolated point, *v*_0_ appears first. Note that each isolated vertex creates a new component. **(C)** At λ = −0.52 points in the denser region appears in the filtration, introducing more components. The indices of the vertices denote the order in which they appear in the filtration. **(D)** At λ = −0.50, all vertices appear in the filtration. Note that, the way we have chosen the filtration function *f*, vertices appear before the edges since *f_v_*(*v*) is always negative. **(E)** At λ = 0.65, the first edge *e*_1_ appears merging two components. By persistence algorithm [8], we pair the edge *e*_1_ with *v*_9_, since *v*_9_ appears later in the filtration. Corresponding to this, we get a persistence pair (*f_v_*(*v*_9_), *f_e_*(*e*_1_)) = (−0.50, 0.65). **(F)** At λ = 0.84, the green edge *e*_2_ appears and creates a cycle. Since there is no 2-simplex(triangle) present, the cycle is never destroyed. In the persistence diagram we have this pair as (*f_e_*(*e*_3_), ∞) = (0.84, ∞). **(G)** At λ = 2.07, the long edge *e*_3_ appears joining *v*_0_ and *v*_1_, yielding a persistence pair (−1.06, 2.07).

Below we briefly describe how we adapt the above persistence framework for analyzing point cloud data (PCD) representing CD8+ T cells in SARS-CoV-2 infection. Details regarding the approach are provided in section 4, the supplementary Algorithms, and Fig S1.

#### Computing persistent homology for cytometry datasets

Our datasets consist of cytometry data for non-naïve CD8+ T cells. Given protein expressions (real values) for *d* proteins in such a single cell, we can represent it as a *d*-dimensional point in 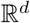. Considering a population of single cells, we get a point cloud (PCD) in 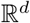. Now, we study the shape of this PCD using the persistence framework that we describe above. We compute persistence diagrams for the PCDs generated with protein expressions from different individuals and compare them. It turns out that, for computational purposes, we need a limit on the dimension *d* for PCD which means we need to choose carefully the proteins that differentiate effectively the subjects of our interest, namely the healthy individuals, COVID-19 patients, and recovered patients. We typically choose 3 (sometimes 2) protein expressions to generate the PCD and call it a PCD in the *P1, P2, P3 space* if it is generated by proteins P1, P2, and P3 respectively.

Flow cytometry data for non-naïve CD8+ T cells in Mathew et al. [3] show generation of CD8+ T cells with larger abundances of the proteins CD38 and HLA-DR (CD38+HLA-DR+ cells) for some COVID-19 patients, forming an “elbow” in the two dimensional PCD with CD38 and HLA-DR protein expressions (see Fig S2). Moreover, there is an increase in the local density of the points (or single CD8+ T cells) in the “elbow” region. This suggests that, to study the PCD generated by the protein expressions by persistence framework, we need to choose a filtration that is able to capture such geometric shapes and variations in the local density.

We briefly describe our choice of filtration by considering the example of a point cloud 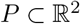 shown in Fig 2. Mathematical and computational details regarding the filtration are provided in the section 4. We build a filtration according to assigned values to the vertices and edges of a graph connecting the input points. For a vertex *p* which is a point in the input PCD *P*, we denote this value *f_v_*(*p*) (given by Eq 1 in Section 4). Similarly, we denote the assigned value to an edge *e* as *f_e_*(*e*) (given by Eq 2 in Section 4); see Fig 2. The values satisfy the conditions that *f_v_*(*p*) < 0 and *f_e_*(*e*) ≥ 0; implications of this specific choice will become clear in the next paragraph. It is noteworthy to mention that *f_v_*(*p*) is the distance-to-measure defined in [17] and captures the density distribution of points whereas *f_e_*(*e*) captures the inter-point distances between the points in the given point cloud.

The persistence algorithm processes each vertex and edge in the order of their appearance in the filtration. We execute it using a threshold value λ from −∞ to ∞ and generate the persistence diagram accordingly. Intuitively, as λ is increased from −∞ to ∞, vertices *p* for which *f_v_*(*p*) ≤ λ and edges for which *f_e_*(*e*) ≤ λ appear in the filtration for a particular value of λ (see Fig 2). Since *f_v_*(*p*) < 0 and *f_e_*(*e*) ≥ 0, all the vertices first appear as λ is increased from −∞ to 0, and then edges start appearing as λ becomes positive. The birth-death events for H_0_ and H_1_ constituting the persistence diagram (Fig 2) contain information about the density and length scales present in the point-cloud. For example, the points showing birth and death events for the H_0_-persistence diagram are more densely organized for the single cell protein expression data from the healthy donor than the SARS-CoV-2 infected patient in the HLA-DR - CD38 plane shown in Fig S3. The denser organization of the birth-death events in the persistence diagram indicates a more homogeneous distribution of CD38 and HLA-DR proteins in the CD8+ T cells in healthy donors compared to that in infected patients. Most of the CD8+ T cells in healthy controls have low amounts of CD38 and HLA-DR abundances and few contain larger values of these proteins, indicating a greater degree of homogeneity (see SI for further details/explanation). The birth-death events for H_1_ in the persistence diagram (Fig S3) in general contain information about the length scales of cyclic structures in the point cloud. It also can capture protrusions like ‘elbows’ that we have in COVID-19 data. Our filtration allows only birth (and not death) of 1-cycles and therefore, a λ value corresponding to the birth of a 1-cycle captures the length scale of the newly born cycle and hence an ‘elbow’. Our analysis of the PCDs in Figure S3D, S3H indeed shows that λ values for the birth of cycles for the COVID-19 patient is much larger compared to that for the healthy individual indicating the presence of larger length scales in the PCD which is consistent with the presence of an “elbow” shape in the PCD for the patient.

### 2.2 Application of persistence to healthy and patient data

Our aim is to find out systematic differences in topological features extracted from cytometry data for healthy individuals and COVID-19 patients. Ideally one would like to compute persistence diagrams for all 25 proteins that were measured in single CD8+ T cells, however, this task encounters two major problems. First, as we mentioned before taking the full 25 dimensional PCD introduces the *curse of dimensionality* [18] making it computationally infeasible to produce the persistence diagrams. The second one is more subtle. In order to measure how the density of data differs from a healthy to infected person in a quantitative way, we need to ensure that the number of points in each PCD, to be analyzed by persistent homology, is the same. Cytometry data usually contain different numbers of single cells in datasets obtained from different donors or replicates. To address the *curse of dimensionality* we use a classifier (XGBoost) that distinguishes single CD8+ T cells in healthy donors from those in COVID-19 patients and we choose the top *r* (taken to be 3) features (proteins) that are deemed important by the classifier while classifying the data points (cells). This reduces the dimension of the data from 25 to a much smaller value denoted *r*. To address the second issue, we perform Kernel Density Estimation (KDE) on every *r*-dimensional dataset and take equal number of samples from it. We then use the filtration defined in Eq 1 & 2 to construct persistence diagrams for each dataset. To quantify the structural differences in the datasets as captured by the corresponding persistence diagrams, we compute the Wasserstein distance [19] between persistence diagrams from randomly selected pairs of either two healthy donors (H×H) or a healthy donor and an infected patient (H×P) and compute distributions of the Wasserstein distances for a large number of (H×H) and (H×P) pairs. The comparison of these distributions provides information regarding the systematic differences in shape features in the CD8+ T cell cytometry data across healthy individuals and COVID-19 patients. The computational pipeline is summarized in (Fig 3). Below we describe results from the application of our computational pipeline to the CD8+ T cell cytometry data in Mathew et al. [3]

**Figure 3.**
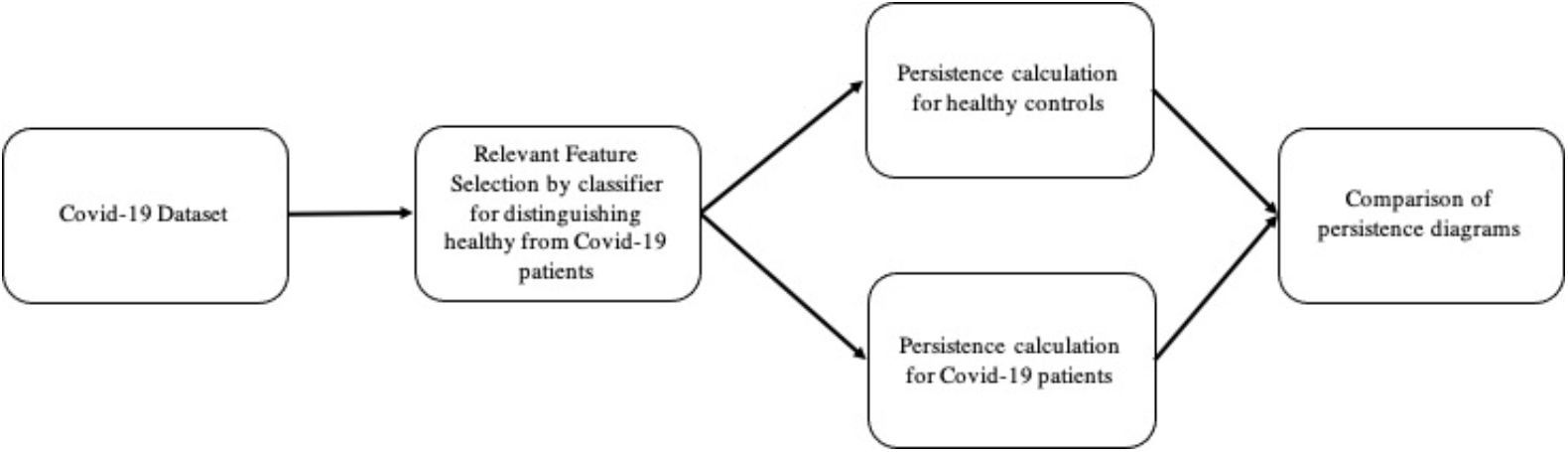
Flowchart of computation pipeline. The pipeline includes three main stages, namely, (i) relevant feature selection, (ii) persistence computation, and (iii) comparison of persistence diagrams.

#### A few protein expressions in CD8+ T cells separate healthy donors from COVID-19 patients

We use the XGBoost, a decision tree based classifier, to rank order proteins for their ability to distinguish CD8+ T cell point cloud data between healthy individuals and COVID-19 patients. The average accuracy of the classifier is about 92%. The classifier returns a feature score for each protein that characterizes its importance relative to other proteins in distinguishing cells from healthy individuals and COVID-19 patients. Intuitively, feature score is an indicator of the importance of a particular feature while classifying the data. By ranking the proteins by their feature scores, we can reduce our further analysis to only a subset of the most important proteins. Our analysis (Fig 4) shows that the top three most important proteins to the XGBoost classifier are proteins T-bet, Eomes, and Ki-67. T-bet induces gene expressions leading to an increase in cytotoxic functions of CD8+ T cells. CD8+ T cells with increased cytotoxic functions are known as “effector” CD8+ T cells and these cells show higher T-bet abundances. Conversely, Eomes induces gene expressions that contribute towards increased life span and re-activation potential of CD8+ T cells to specific antigens [20]. These long-lived T cells are known as “memory” T cells which show increased expressions of Eomes. Memory T cells provide key protection against re-exposure to the same infection. Ki-67 is a marker for actively proliferating cells [21]. Mathew et al. [3] identified Ki-67 as one marker that is upregulated (increased) in some COVID-19 patients. These three proteins are most likely to distinguish CD8+ T cells in healthy donors from those in patients. Further details regarding the application of the classifier are provided in the Materials and Methods section (section 4).

**Figure 4.**
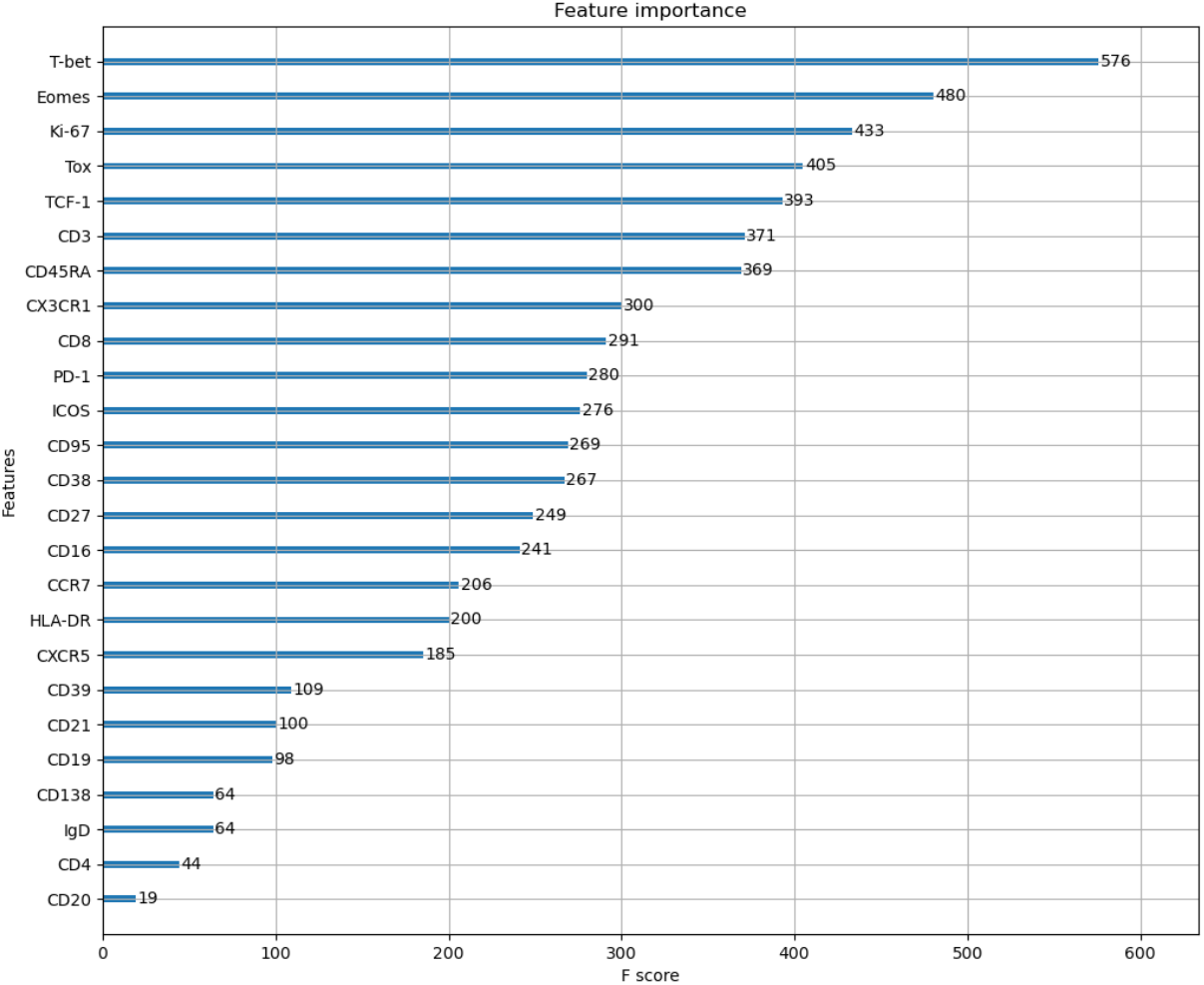
Rank ordering of proteins using a decision tree based classifier. Shows rank ordering of proteins by descending values of feature importance generated by the classifier XBoost.

#### Persistence diagrams distinguish structural features in CD8+ T cell data occurring in healthy individuals and COVID-19 patients across batch effects and donor-donor variations

We select the proteins T-bet, Eomes, and Ki-67 as relevant markers and compute the persistence diagrams of the PCD given by them for each individual belonging to groups of healthy donors, COVID-19 patients, and recovered patients. The persistence diagrams vary from individual to individual in each group and between groups which could arise due to batch effects in samples and/or donor-to-donor variations. To determine if there are systematic differences in persistence diagrams for individuals across the three groups (healthy, patient, and recovered), we compute Wasserstein distance between persistence diagrams for 3 categories of pairings: 1) two healthy donors (H×H), 2) one healthy donor and one patient (H×P), and 3) one healthy donor and one recovered individual (H×R). We compute distances for 108 randomly chosen pairs of individuals for each category of pairings. Wasserstein distances of the persistence diagrams for 0-th and 1-st homology groups H_0_ and H_1_ respectively are higher when comparing H×P pairs than when comparing Hx H pairs (Fig 5). This indicates that systematic geometric differences in the flow cytometry PCD with T-bet, Eomes, and Ki-67 between individuals with and without COVID-19 are not attributable to batch effects or donor-to-donor variations alone. Increasing the number of randomly chosen pairs to 200 did not change the qualitative differences shown in Fig 5 (Fig S4). The difference between H×H and H×R distributions of distances in the T-bet, Eomes, and Ki-67 space are less prominent (Fig S5). We further test if such systematic differences are present for proteins that are at the bottom of the list in Fig 4 and find that the distributions of Wasserstein distances for corresponding persistence diagrams overlap between the H×H and H×P pairs (Fig S6). This suggests that systematic differences in the geometry of the PCD occur only for specific sets of proteins. Details regarding computation of persistence diagrams and Wasserstein distances are given in Section 4.

**Figure 5.**
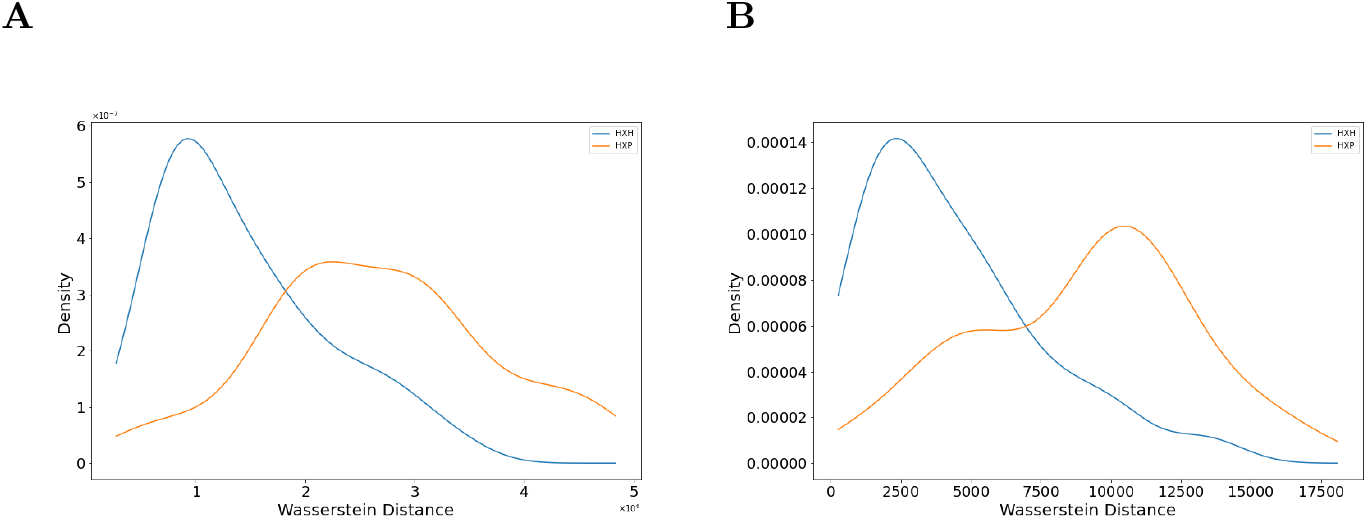
Distributions of Wasserstein distances between persistence diagrams. **(A)** Shows distributions of Wasserstein distance between H_0_-persistence diagrams for H×H (blue line) and H×P (orange line)pairs. **(B)** Shows distributions of Wasserstein distance between H_1_-persistence diagrams for H×H (blue line) and H×P (orange line) for the same pairs in (A).

Next, we select a comparison pair that generates a large Wasserstein distance between H_1_-persistence diagrams to further investigate what structural differences exist between the datasets. We choose one pair of a healthy control and patient that generated a Wasserstein distance of 4.0 × 10^6^ units in their H_0_-persistence diagrams and 1.1 × 10^4^ units in H_1_-persistence diagrams. These two individual PCDs and their resulting persistence diagrams are shown in Fig 6.

**Figure 6.**
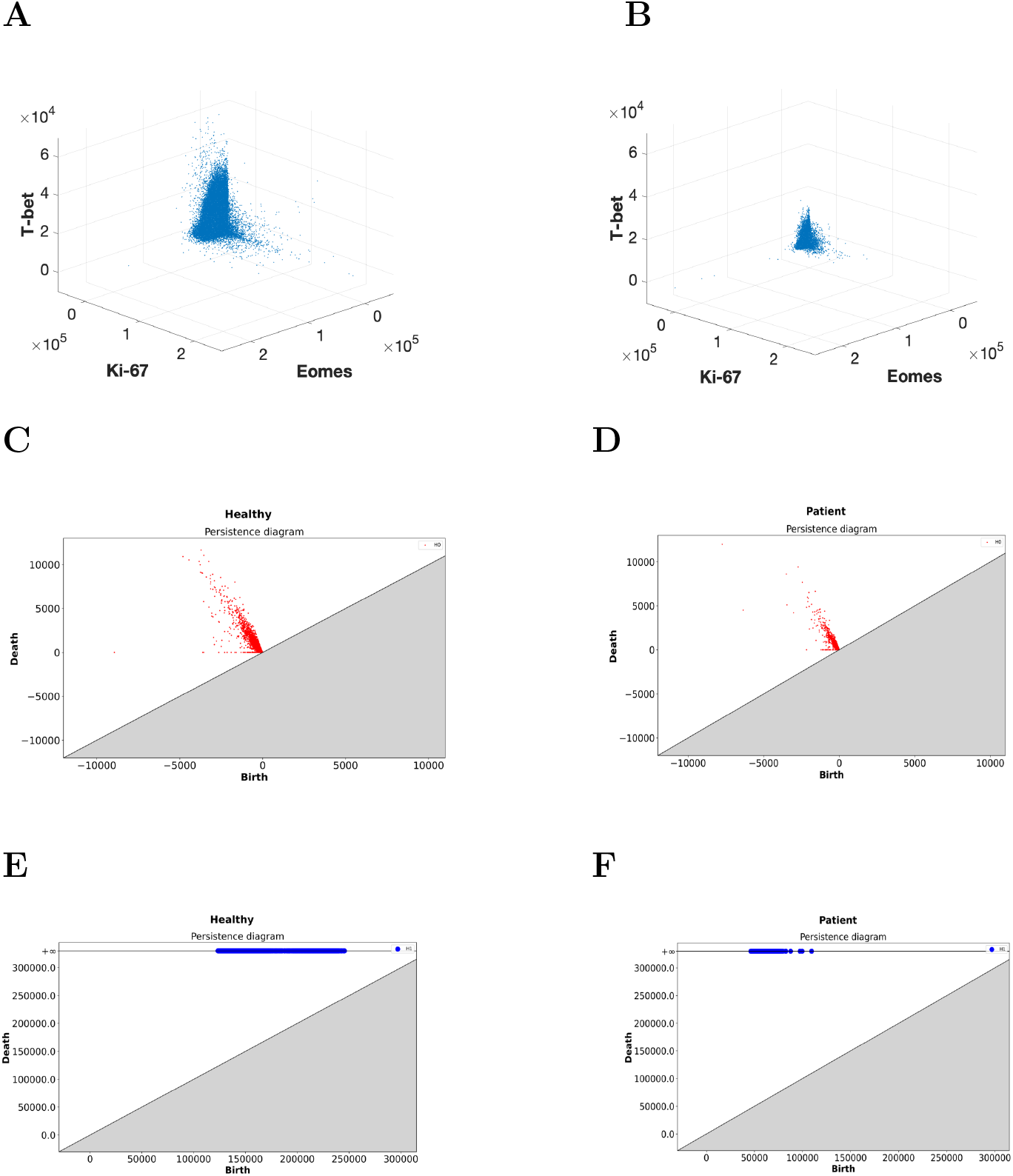
Differences in shape features in the 3D point cloud for CD8+ T cells in a H×P pair. CD8+ T cell point cloud for proteins Eomes, Ki-67, and T-bet for **(A)** a healthy control and **(B)** a COVID-19 patient. **(C)** Shows H_0_-persistence diagram for the healthy control in (A). **(D)** Shows the H_0_-persistence diagram for the COVID-19 patient in (B). **(E)** H_1_-persistence diagram for the healthy control in (A). **(F)** H_1_-persistence diagram for the COVID-19 patient in (B).

A readily apparent difference between the resulting persistence diagrams is given by the lower birth times in H_1_ of the COVID-19 patient compared to the healthy control (Fig 6e, 6f). This result indicates that the length scale of the data is smaller in the COVID-19 patient, which can be visually confirmed in the scatter plots of the data (Fig 6a, 6b). Specifically, the single cell abundances of T-bet and Eomes in CD8+ T cells are clustered significantly tighter around the origin for the COVID-19 patients than for the healthy controls. Similar manual inspection of other H×P pairs that generate large Wasserstein distances between their persistence diagrams confirms that this trend is not limited to this pair alone.

Additionally, the points in the H_0_-persistence diagram are spread out more widely for the healthy control than the COVID-19 patient (Fig 6c, 6d). A wider distribution of births and deaths in the 0-th homology H_0_ implies that there are regions of disparate densities. This suggests that the densities in the protein expressions of T-bet and Eomes are more homogeneous in the PCD in the COVID-19 patient than in the healthy control.

The structural change in the PCD for CD8+ T cells in the T-bet/Eomes plane that occurs during COVID-19 infection implies that T-bet and Eomes expression should be downregulated (decreased) in non-naïve CD8+ T cells. This result is consistent with analysis of clusters of CD8+ T cells by Mathew et al. [3] that shows that clusters high in T-bet and/or Eomes are downregulated in COVID-19 patients. The downregulation of T-bet and Eomes in response to viral infections is not well documented, as CD8+ T cells commonly differentiate into phenotypes with, high T-bet, high Eomes, or both in response to infections [20, 22].

## 3 Discussions and conclusions

We developed a persistent homology based approach to determine topological features hidden in point cloud data representing single cell protein abundances measured in cytometry data. In particular, we characterized the number of connected components or H_0_, and the number of holes or H_1_ in our persistence calculations, and showed that our approach is able to determine systematic shape differences in the cytometry data for CD8+ T cells obtained from healthy individuals and COVID-19 patients. Therefore, the approach is able to successfully determine systematic shape differences that exist in the presence of batch effect noise and donor-donor variations in the cytometry data. Furthermore, our approach does not use data transformations (e.g., arc-sinh transformation) or any ad-hoc subtype gating to determine these systematic differences, thus we expect persistent homology based approaches will be especially useful in identifying high-dimensional structural trends hidden in cytometry data.

We determined structural changes in T-bet and Eomes abundances in single CD8+ T cells in COVID-19 patients that can be summarized as downregulation. This result is non-intuitive as previous findings show that T-bet and Eomes protein abundances are highest in effector CD8+ T cells which are induced in response to acute infections suggesting T-bet and Eomes expressions are upregulated in CD8+ T cells responding to infections [20, 22]. The clinical implications of this result are unclear, Mathew et al. [3] describe a immunophenotype in which Eomes+, T-bet+, CD8+ T cells are more abundant in COVID-19 patients who respond poorly to Remdesevir and NSAIDs, have high levels of IL-6, and have fewer eosinophils. Our analysis identifies that this immunophenotype (i.e.,Eomes+, T-bet+, CD8+ T cells) is systematically less prevalent in COVID-19 patients than in healthy controls. The ability of our approach to identify non-intuitive shape features such as the above without any ’supervision’ (e.g., specific gating) of the cytometry data shows that it can potentially determine more complicated immunologically relevant shape features. For instance, although we did not identify differing shapes or voids in the COVID-19 data in our current approach, our method would be able to detect these in a different dataset that contains such characteristics.

Our approach integrates cellular comparisons with dataset comparisons. First, the classifier pools all data and determines which proteins are significant in discriminating whether cells come from healthy controls or COVID-19 patients. In this way, the classifier identifies a way to compare cellular phenotypes across experimental groups. Next, the computation of Wasserstein distances for persistence diagrams compares individuals against each other, integrating cellular phenotypes with donor information (e.g., healthy and COVID-19 patients). Thus, this approach allows us to automatically identify individuals that are associated with distinguishing structural features in the point cloud data.

Currently, the limitations are mostly centered around computational complexity and the curse of dimensionality. The computational resources necessary to calculate a KDE are great enough that extensions beyond three dimensions are not feasible. However, improving or circumventing the KDE calculation step will significantly increase the dimensionality of data that can be considered. Since we are computing pairwise distances between datapoints to obtain the persistence diagram (Section SIB), computation time will linearly increase as the dimension of data increases. Our methodology on our computational cluster resources currently takes about 40 minutes. This is comparable to other data science applications to large datasets, but can be a barrier to those without access or experience with computational clusters. Additionally, it is unclear how adding more dimensions will impact the statistical properties of the data and interpretability of the results. To expand into many (i.e. 25) dimensions, computational interpretation and validation tools will be necessary.

## 4 Materials and Methods

### Relevant feature selection by the XGBoost classifier

Let *D* = (*c*_1_, *c*_2_, …, *c_m_*} be the collection of *m* cytometry datasets. Each dataset, *c_i_*, can be viewed as a 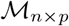 matrix where *n* is the number of datapoints (cells) and *p* is the number of proteins with which each *c_i_* is generated. We denote the collection of cytometry datasets of healthy individuals as 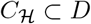 and similarly the set of individuals infected with SARS-CoV-2 as 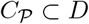. We proceed to label the data in the following manner: If 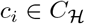 then we assign the label +1 to each of the *n* datapoints, similarly we assign −1 if 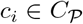. Essentially, we now have a binary classification problem where our labeled dataset is 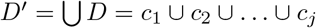, with labels defined as above. We solve this binary classification problem with XGBoost [23], a gradient boosted decision tree based classifier, and as a byproduct we get feature scores that correspond directly to each feature’s importance in the classification. The higher the score for a protein, the more important it is for the classifier’s decision. After our classifier orders the proteins by their scores, we take first *r* proteins to construct the point-cloud on which persistence diagrams are computed. We set *r* = 3 for all our analysis reported here. We used data from 56 healthy individuals and 108 COVID-19 patients for our feature selection.

The XGBoost classifier was implemented using the open-source python XGBoost package [23]. The model was then trained and validated with *K*-fold cross-validation, with *K* =10. The average accuracy of the classifier was 92.14 ± 0.04%. The protein scores are shown in Fig 4.

### Kernel Density Estimation

We estimated the probability distribution function associated with the point cloud data in three dimensions using Kernel Density Estimation. The estimated probability distribution function was used to draw 20,000 points corresponding to each PCD that were further analyzed using persistent homology. We used KDE estimated probability distributions instead of random selection of 20,000 datapoints in three dimensions, because the random selection of data points do not appropriately sample the distribution of the data points for the relatively smaller sample size (20, 000 data points). KDE was implemented using scikit-learn [24] package. The kernel function was chosen to be ‘gaussian’. For now we have tuned the bandwidth parameter based on manual observation and to remain consistent we have fixed the bandwidth at 0.2 for each dataset.

### Details of persistent homology computation

As mentioned before (section 2.1), computation of persistence diagrams needs a *filtration*. We set the filtration induced by the function *f* = {*f_v_*, *f_e_*} where *f_v_*(*p*) computes an “average” Euclidean distance between the vertex *p* and its *k* neighbors according to Eq. (1) and *f_e_*(*e*) computes the length of the edge *e* according to Eq. (2).

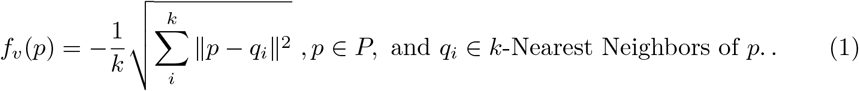

The term ‖*p* − *q_i_*‖ in the above equation is the Euclidean distance between the vertices *p* and *q_i_*. The function value *f_e_*(*e*) for an edge *e* = (*p, q*) is given by the Euclidean distance between *p* and *q*. For the experiments, the number of nearest neighbors is fixed to *k* = 40.

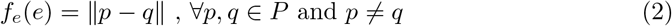

We begin with computing Kernel Density Estimation (KDE) for every cytometry PCD *c_i_* and take *n*(= 20,000) samples from each KDE. We do this to make *c_i_* uniform *w.r.t*. number of data points (single CD8+ T cells). We compute a complete weighted graph *G*(*V, E*) with vertices in the sampled data. This complete graph *G* is a key-step that enables us to compute the *persistence diagram, Dgm*(*c_i_*) of the dataset *c_i_*, *w.r.t*. the filtration function *f*. We show the algorithm (Algorithm 2) that executes this step in detail in the supplementary material. Notice that the graph *G* is weighted in the sense that each vertex *v* ∈ *V* and edge *e* ∈ *E* carries a weight of *f_v_*(*v*) and *f_e_*(*e*) respectively. Observe that 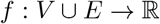 constitutes a valid filtration of *G*.

We compute persistence diagrams for each *c_i_* ∈ *D* according to Algorithm 3. The next step involves comparing the persistence diagrams. We do this by computing the Wasserstein distance between persistence diagrams and plotting their distributions. We take two persistence diagrams of randomly selected healthy individuals and compute the Wasserstein distance between them with the help of Gudhi [19, 25] and scikit-learn Python library [24]. Similarly, we compute Wasserstein distance between persistence diagrams of a healthy and an infected individual (both are randomly drawn from the collection). We plot the resulting distances. We do this for 108 pairs to obtain two distributions. Note that, results described in Section 2.1 still holds for 200 pairs (Fig S4). Intuitively, a large Wasserstein distance between two persistence diagrams implies the datasets on which they were constructed are structurally very different while a small distance implies they are structurally similar.

### Flow cytometry data for healthy individuals and COVID-19 patients

The data come from Mathew et al., 2020 [3] and was retrieved from Cytobank. Mathew et al. performed high-dimensional flow cytometry experiments using peripheral blood obtained from 125 patients admitted to the hospital with COVID-19, 36 donors that recovered from documented SARS-CoV-2 infection, and 60 healthy controls. Our analysis focuses on the deposited data available at https://premium.cytobank.org/cytobank/experiments/308357 for non-naïve CD8+ T cells collected at the time of admission (and not any later blood draws, such as at 7 days after admission). We removed forward- and side-scatter variables and other non-protein measurements, resulting in 25 proteins included in our analysis.

### Available code

Our current code is available at https://github.com/soham0209/TopoCytometry and will be updated for ease-of-use and performance enhancements.

## Acknowledgements

This work is supported by the NIH awards R01-AI 143740 and R01-AI 146581 to JD, and by the Research Institute at the Nationwide Children’s Hospital, and NSF awards CCF 1839252 and 2049010 to TD. The authors would like to thank John Wherry and members of his lab for depositing their data and helping us to understand it.

