## Supplementary Materials for "Determining clinically relevant features in cytometry data using persistent homology"

### A Some omitted details

We have already introduced the concept of persistent homology. Its complete exposition is beyond the scope of this paper. We provide another intuitive example to illustrate its application in capturing prominent features hidden in a PCD. We refer to [8, 26] for a detailed exposition of topological persistence.

Suppose a set of points $P$ is sampled along a curve with two 'holes'; see Figure S1. If we grow balls of radius $\epsilon$ starting from zero around the sampled points we see that different holes get filled up at different times. The bigger hole gets filled at a much later time than the small one and the spurious ones. Persistent homology formalizes this idea of tracking the lifetime of topological features (homology groups).

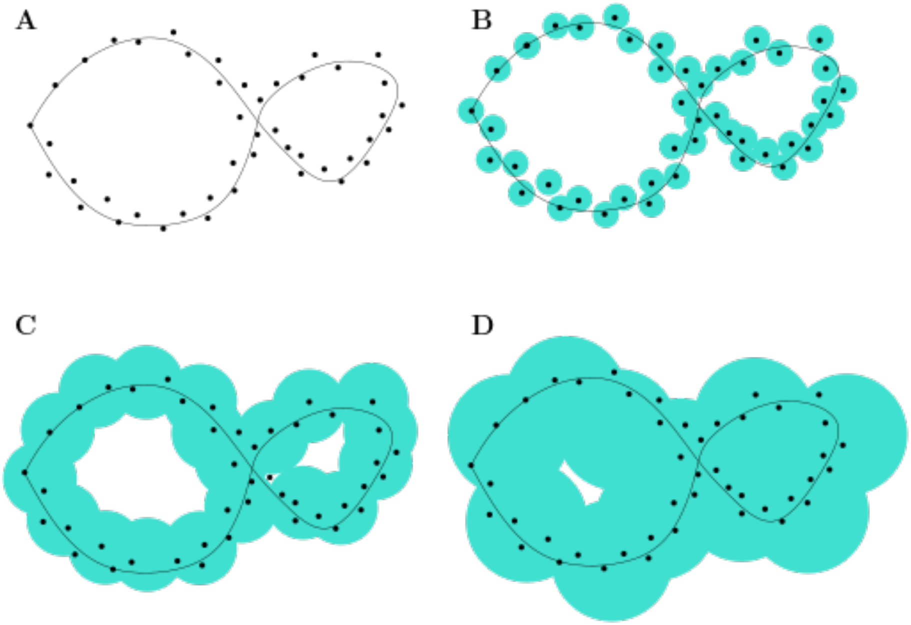


**Figure S1. (A)-(D)** Shows the concept of persistence intuitively. **(A)** A set of points $P$ sampled from a curve. **(B)** An Euclidean ball of radius $\epsilon$ is grown around each point in $P$. **(C)** As $\epsilon$ increases the smaller hole gets filled up. **(D)** The larger hole still 'persists' even though the smaller hole gets filled. Figures are adopted from [26, Fig 4.2].

**Definition A.1** (Simplex)**.** A $k$-simplex $\sigma$ is the convex hull of $k+1$ affinely independent point set $P$. We call $k$ to be the dimension of $\sigma$ and denote $dim(\sigma) = k$. A *face* $\sigma'$ of $\sigma$ is the convex hull of non-empty subset of $P$ and this relation is given by $\sigma' \subseteq \sigma$.

In particular 0-simplex is a point, 1-simplex is an edge, 2-simplex is a triangle and so on.

**Definition A.2** (Simplicial Complex)**.** A set of simplices is defined as a simplicial complex $\mathcal{K}$ if the following two restrictions hold

- If $\sigma \in \mathcal{K}$ and $\sigma' \subseteq \sigma$ then $\sigma' \in \mathcal{K}$.
- For any two simplices $\sigma, \tau \in \mathcal{K}$, $\sigma \cap \tau$ is either empty or a face of both $\sigma$ and $\tau$.

The *dimension* of a simplicial complex $\mathcal{K}$ is the maximum dimension of any of its simplices.

**Algorithm 1** DisPers

**Input:** $c$: Cytometry dataset, $k$: Nearest neighbors to consider, $n$: Number of samples
**Output:** $Dgm(c')$: Persistence diagram of $c'$ sampled from cytometry data $c$

**begin**
  Compute KDE on $c$
  $c' \leftarrow$ Generate $n$ samples from KDE on $c$
  $G \leftarrow$ GEN-COMPLETE-GRAPH($c'$, $k$)
  $Dgm(c') \leftarrow$ COMPUTE-PERSISTENCE($G$)
  **return** $Dgm(c')$
**end**

**Definition A.3** (Filtration)**.** Given a Simplicial Complex $\mathcal{K}$ and a *monotonic* function $f : \mathcal{K} \to \mathbb{R}$ we get a nested sequence of subcomplex by denoting $\mathcal{K}_i = f^{-1}(-\infty, a_i]$ which is defined as *filtration*:

$$\phi = \mathcal{K}_0 \subseteq \mathcal{K}_1 \subseteq \mathcal{K}_2 \subseteq \ldots \subseteq \mathcal{K}_n = \mathcal{K}$$

By *monotonic* we mean that if $\sigma' \subseteq \sigma$, $f(\sigma') \leq f(\sigma)$.

### B Algorithms

Since a graph is a 1-dimensional simplicial complex, for a graph with $m$ edges and $n$ vertices, we can compute its 0-dim persistence in $\mathcal{O}(m \log n)$ time with Kruskal like minimum spanning tree algorithm [8].

**Algorithm 2** Gen-Complete-Graph

**Input:** $c$: Sampled cytometry data, $k$: Nearest neighbors to consider
**Output:** $G(V, E)$: Complete Weighted Graph on $c$

**procedure** GEN-COMPLETE-GRAPH
  $G(V, E) \leftarrow \phi$  ▷ $V$ is the set of nodes and $E$ is the set of edges of $G$
  **for all** $v \in c$ **do**
    Compute $v_1, v_2, \ldots, v_k$, k-nearest neighbors of $v$
    $w(v) \leftarrow -\frac{1}{k}\sqrt{\sum_i \|v - v_i\|^2}$  ▷ $w(v)$ denotes vertex weight
    $V(G) \leftarrow V(G) \bigcup \big(v, w(v)\big)$
  **end for**
  **for all** $\{u, v\} \in c \times c$ and $u \neq v$ **do**
    $e \leftarrow \{u, v\}$
    $w(e) \leftarrow \|u - v\|$  ▷ $w(e)$ denotes weight of the edge
    $E(G) \leftarrow E(G) \bigcup \ (e, w(e))$
  **end for**
  **return** $G(V, E)$
**end procedure**

Algorithm 3 shows the steps of Persistence computation. Consider a generic step when an edge $e \in G$ is introduced in the minimum spanning forest that joins two forests rooted at nodes $v_0$ and $v_1$. For the new tree we will choose the root as node having smaller weight and the edge $e$ pairs with the node having larger weight. Essentially we are choosing to kill the youngest connected component created by, with a little abuse of notation, the vertex $argmax(w(v_0), w(v_1))$. We define persistence of the edge as

$$p(e) = w(e) - \max\{w(v_0), w(v_1)\} \tag{3}$$

It is important to point out that an edge may or may not necessarily kill a connected component. If it does not kill a connected component it definitely creates a 1-cycle and we pair the edge with a special vertex with $w(v) = \infty$ and define $pers(e) = \infty$.

---

**Algorithm 3** Compute Persistence Diagram

---

**Input:** $G(V, E)$: Complete weighted graph on sampled cytometry data $c$
**Output:** $Dgm(c)$: Persistence Diagram

**procedure** COMPUTE-PERSISTENCE
  $E' \leftarrow$ Sort $E$ in increasing order of $w(e)$ with $e \in E$
  $P_0 \leftarrow \phi$ ▷ $P_0$ tracks 0-dim birth and death
  $P_1 \leftarrow \phi$ ▷ $P_1$ tracks 1-dim birth
  **for all** $e = (u, v) \in E'$ **do**
    $Root_0 \leftarrow find(u)$
    $Root_1 \leftarrow find(v)$
    **if** $Root_0 \neq Root_1$ **then**
      $birth \leftarrow max\{w(Root_0), w(Root_1)\}$ ▷ $e$ kills youngest homology class
      $death \leftarrow w(e)$
      $pers(e) \leftarrow death - birth$
      $merge(Root_0, Root_1)$
      $P_0 \leftarrow P_0 \cup (birth, death)$
    **else**
      $P_1 \leftarrow P_1 \cup (w(e), \infty)$ ▷ $e$ is a creator and creates a 1-cycle
    **end if**
  **end for**
  $Dgm(c) \leftarrow \{P_0, P_1\}$
  **return** $Dgm(c)$
**end procedure**

---

### C Supplementary Figures

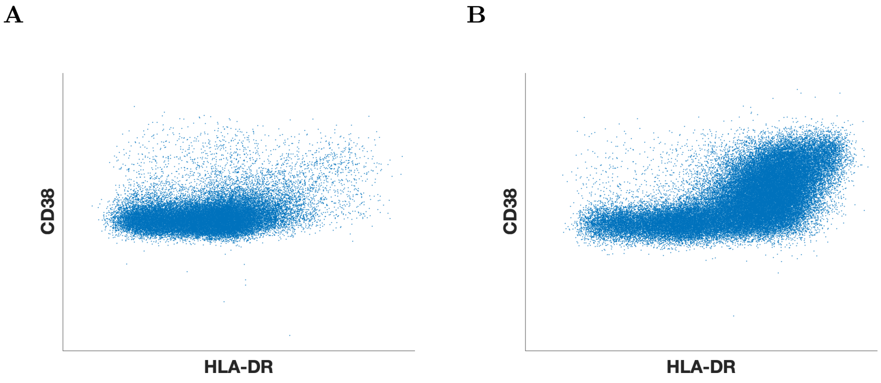


**Figure S2.** Transformed scatter plot for HLA-DR/CD38 axes in a singular **(A)** healthy donor and **(B)** COVID-19 patient. This plot demonstrates the “elbow” found by the authors in [3]. The x-axis is asinh(HLA-DR/200) and the y-axis is asinh(CD38/500)

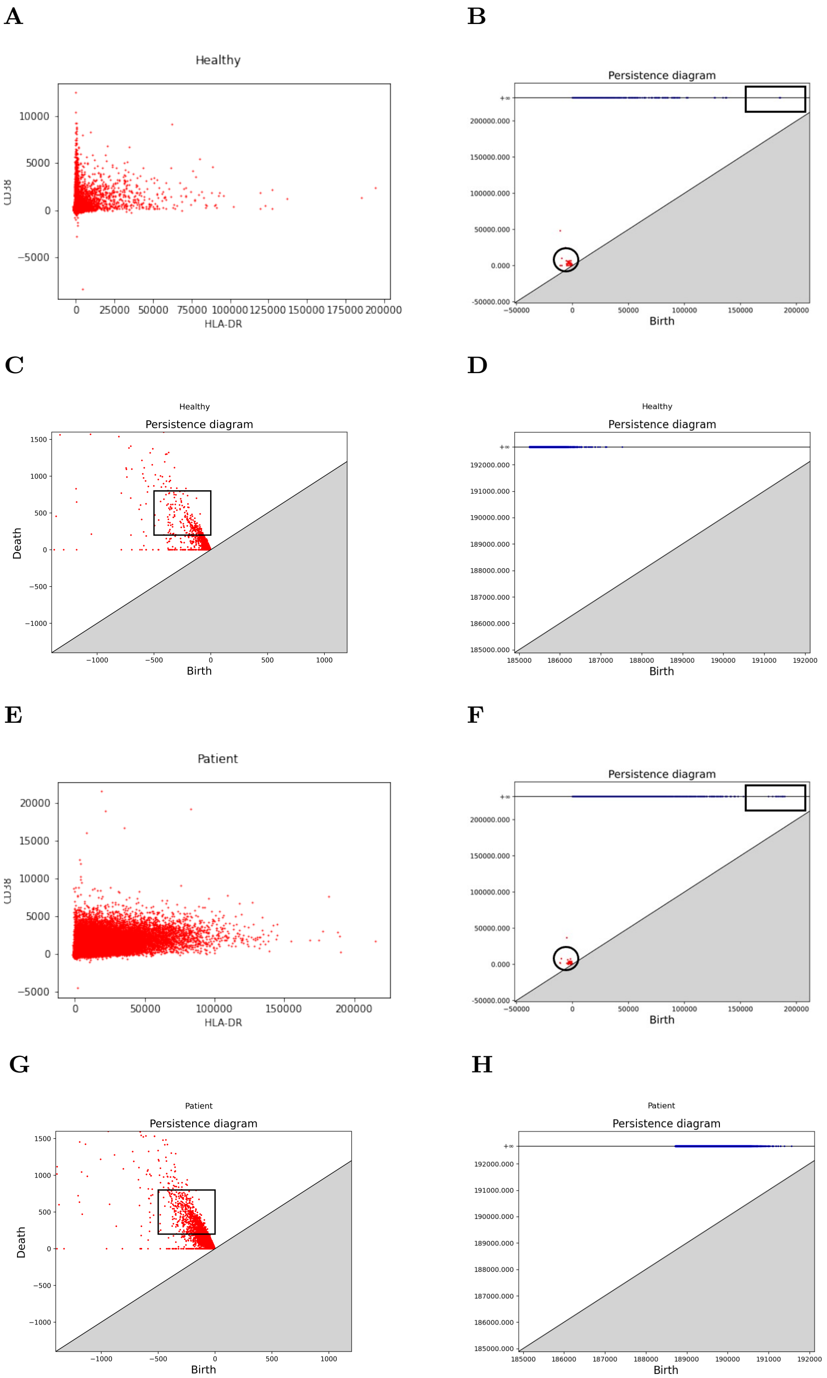


**Figure S3.** Persistence calculations and comparisons for HLA-DR/CD38 axes shown in Mathew et. al. [3] **(A)** Point cloud for individual healthy control in HLA-DR/CD38 expression levels. **(B)** Complete persistence diagram for the healthy control shown in (A). Boxes indicate zoomed regions for figures (C) and (D); **(C)** Zoomed in region from (B) of $\mathsf{H}_0$ persistence diagram. Box shows area of low density compared to patient persistence diagram; **(D)** Zoomed in region from (B) of $\mathsf{H}_1$ persistence diagram; **(E)-(H)** Same as (A)-(D), but for an individual COVID-19 patient. The box in (G) is more densely populated than the identical box in (C).

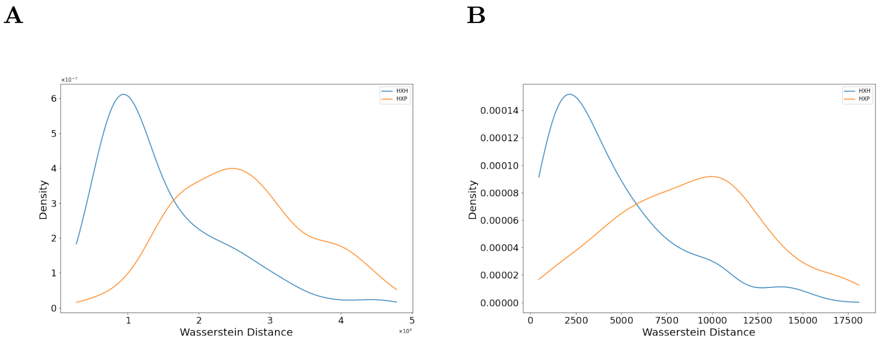


**Figure S4.** Distributions of Wasserstein distances between **(A)** $H_0$-persistence diagrams and **(B)** $H_1$-persistence diagrams. Distances between pairs of healthy controls (H × H) and pairs of a healthy control and a COVID-19 patient (H × P) are overlaid. Persistence diagrams are calculated from point clouds in the T-bet, Eomes, and Ki-67 axes. This figure plots distributions of **200** randomly selected pairs, while Fig 5 plots distributions of 108 randomly selected pairs.

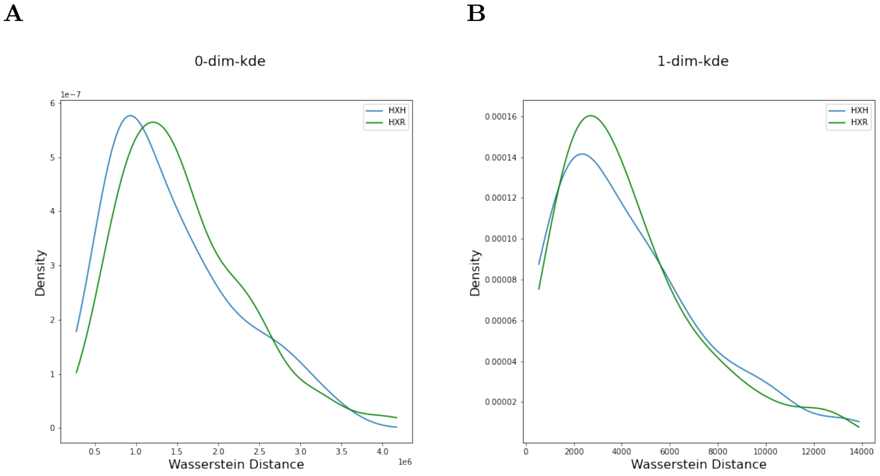


**Figure S5.** Distributions of Wasserstein distances between **(A)** $H_0$-persistence diagrams and **(B)** $H_1$-persistence diagrams. Distances between pairs of healthy controls (H × H) and pairs of a healthy control and a individual that recovered from COVID-19 (H × R) are overlaid. Persistence diagrams are calculated from point clouds in the T-bet, Eomes and Ki-67 axes.

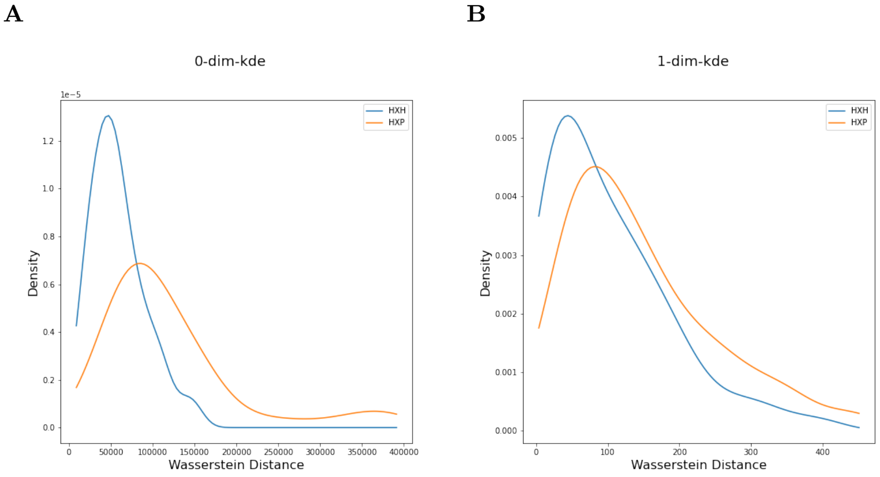


**Figure S6.** Distributions of Wasserstein distances between **(A)** $\mathsf{H}_0$-persistence diagrams and **(B)** $\mathsf{H}_1$-persistence diagrams. Distances between pairs of healthy controls (H $\times$ H) and pairs of a healthy control and a COVID-19 patient (H $\times$ P) are overlaid. Persistence diagrams are calculated from point clouds in the IgD, CD4 and CD20 axes.
